# Recent Urban Development Reduces Bee Abundance and Diversity

**DOI:** 10.1101/2021.11.19.469286

**Authors:** Vera Pfeiffer, David W. Crowder, Janet Silbernagel

**Author notes:** Corresponding author: Vera Pfeiffer.

## Abstract

Wild bee communities persist in cities despite major disruption of nesting and food resources by urban development. Bee diversity and abundance is key for urban agriculture and maintenance of plant diversity, and assessing what aspects of cities enhance bee populations will promote our capacity to retain and provision bee habitat. Here, we assessed how variation in land cover and neighborhood development history affected bee communities in the midwestern US urban landscape of Madison, Wisconsin. We sampled bee communities across 38 sites with relatively high (> 55%) or low (< 30%) levels of impervious surface, and assessed effects of land use and neighborhood development history on bee abundance and species richness. We show abundance and richness of bees was lower in recently developed neighborhoods, with particularly strong negative effects on soil nesting bees. Soil nesting bees and bee community richness decreased as cover of impervious surface increased, but above ground nesting bees were minimally impacted. Bee community similarity varied spatially and based on dissimilar local land cover, only for soil nesting bees, and the overall bee community. Impervious surface limited bee abundance and diversity, but new neighborhoods were associated with greater negative effects. We suggest that enhancing the structural diversity of new neighborhoods in urban ecosystems may imitate the structural benefits of older neighborhoods for bee populations.

## Introduction

Urban development is rapidly transforming the Earth’s surface. Impervious surfaces and fragmented patches of vegetation that typify urban ecosystems threaten species diversity (Rebele 1994). In urban ecosystems, ecological communities are also exposed to loss of food and nesting resources, despite cultivated gardens adding diversity (Rebele 1994, Rosenzweig 2003). Urban habitat fragmentation often leads to the loss of plants and associated pollinators, especially plants reliant on animal pollination (Biesmejer et al. 2006; Theodorou et al. 2020). However, organisms differ in their sensitivity to urbanization, and certain pollinators thrive in urban ecosystems despite high disturbance. More research is thus needed to assess relationships between ecological community structure and land use in urban landscapes to protect biodiversity and ecosystem services in ecosystems supporting most of the human population (Daily et al. 1997).

Native bees promote diversity of urban ecosystems by pollinating native, ornamental, and agricultural plants across seasons (Hoehn 2008, Garibaldi 2013). As urban development expands, urban agriculture is growing, emphasizing the need to maintain urban pollinators to produce food where people live (Hodgson et al. 2011). Habitat simplification and high density of honey bee apiaries in urban systems can negatively affect wild bees (Gonzales et al. 2013, Martins et al. 2013; Renner et al. 2021), but high bee diversity has also been observed in cities like New York and Chicago, US (McFrederick and LeBuhn 2006; Matteson et al. 2008; Fetridge et al. 2008). This shows urban land can provide diverse floral resources, especially when gardens provide flowers for a longer duration than other ecosystems (Goddard et al 2010, Threlfall et al. 2015). Urbanization may have different impacts on bees with different ecology, however.

Within urban ecosystems, variation in pollinator nesting strategy may predict sensitivity of species to the high levels of disturbance in urban systems. In many cases, below-ground nesting cavity bees are expected to be more affected by urbanization than bees that nest aboveground given the prevalence of impervious surfaces (Larsen 2005, Cane et al. 2006; Jha and Kremen 2013). For example, many bees excavate or construct their own nesting cavities using mud, wood or pithy stems, or dig cavities in the soil, and these habitats are often less available in urban compared to natural or rural landscapes. However, man-made structures can in some cases supplement nesting habitat, by providing stone walls, wooden structures, and various other cavities, as well as bare ground and loosened soil. By investigating what aspects of land cover and land use underlie trends in species filtering, we can increase our capacity to restore the resources that are lost along with associated taxa.

Here we assessed effects of land cover and neighborhood development on the urban bee communities associated with the growing urban cityscape of Madison, Wisconsin, United States. Our study tested three main hypotheses. First, we predicted that property development would increase the amount of impervious surface area, disturbing bee habitat and reducing abundance and species richness of bee communities. In particular, we expected stronger effects of property development on below-ground cavity nesting bees that require already excavated cavity spaces, often underground. Second, we predicted that bee community composition would be more dissimilar with greater geographic distance across the city, especially for small soil-nesting bees with limited dispersal capacity. Third, we predicted that property development would decrease similarity in the composition of bee communities. By assessing effects of land cover, property development, and spatial scale on species richness and species composition of bee communities, our study contributes to the empirical foundations of pollination ecology as it relates to conservation and restoration efforts in urban ecosystems.

## Materials and methods

### Study area and sampling design

Madison, Wisconsin is an urban state capital surrounded by agricultural land in one of the fastest growing counties in the US. The primary transition type occurring in the Madison area for the past century is the conversion of agricultural to urban land around the city edge (Wegener 2001; Carpenter et al. 2007; Riera et al. 2001). The dominant urban area is typified by mixed residential and commercial zones with small forest patches and city parks. The 123 km^2^ central urban zone of Madison includes 46 km^2^ (37%) of impervious surface, 30 km^2^ (24%) of vegetated space, with the remaining landscape covered by lakes. The city receives semi-frequent rain and severe thunderstorms throughout the summer months that supports abundant flowering prairie plants in city parks or where native grasslands have been conserved or restored around the city.

Flower-visiting insects were sampled across Madison using a spatially stratified survey to account for changing regional species pools. To select sites, a grid of 2.5 2.5km squares was laid across Madison and cells dominated by lake or agriculture were excluded, leaving 19 cells dominated by high-density residential and urban land (Fig. 1). In each of these cells we used a paired design and selected two sites characterized by either (1) high (> 55%) or (2) low (< 30%) impervious surface area within the surrounding 200 m based on a lower resolution classified land cover surface (USDA-NASS 2013). Within each cell, paired sites with high or low impervious surface area were separated by at least 400 m. These 38 sites were selected in a stratified-random manner, and permission from property owners (identified from a city database) was requested until appropriate locations were identified. Sample sites included primarily residential properties, as well as commercial properties, urban storm water management areas, and city parks.

**Figure 1.**
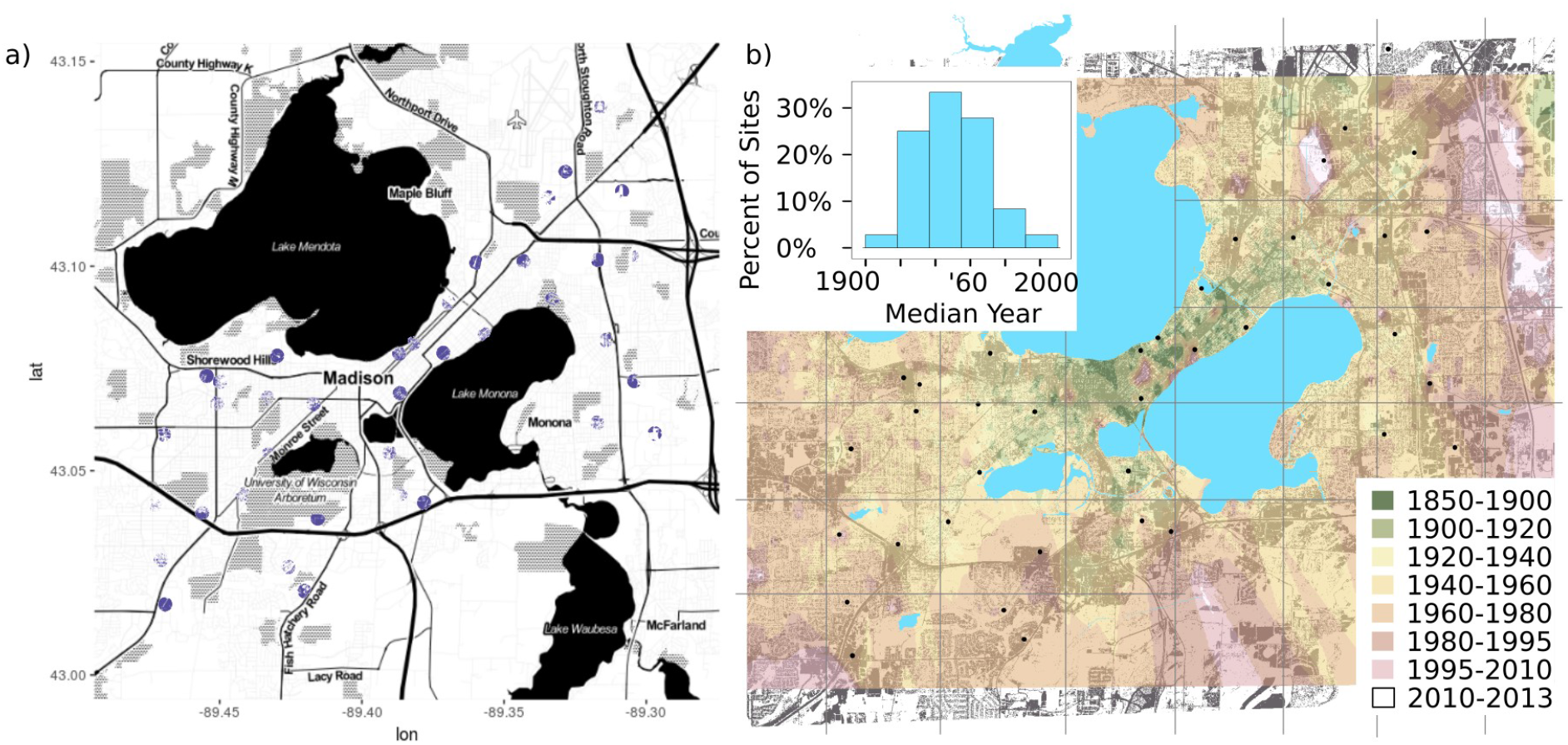
Map of urban bee community sampling sites selected in a spatially stratified design across the City of Madison, (a) surrounded by 200m buffers, filled with impervious land cover and (b) overlaid on kriged property development year surface.

### Bee community sampling

Bees were sampled six times between early June and late August 2013. Pan traps were distributed every two weeks during clear, sunny days when bees were foraging. All traps were distributed across the same evening to early morning period (after 17:00-dark and before dawn-8:00), and collected 4 d later. Six bee traps were placed at least 5 m apart within a 40 m area in each site, with two dark blue, two canary yellow, and two white; bees were also trapped in 0.5 L pan traps suspended 20 cm or 2.5 m from the ground to match the height of flowering vegetation. Bees were identified to species using the discover life online key and a comprehensive dichotomous key available for *Lasioglossum* (Ascher & Pickering 2013; Gibbs 2011).

We classified bee taxa as soil-nesting, below-ground cavity-nesting, and above-ground nesting bees, based on available observations. The below ground cavity-nesting bees included 7 species of bumble bees (*Apidae: Bombus*). Above-ground nesting bees included small carpenter bees (*Ceratina spp*.), yellow faced bees (*Hylaeus spp*.), carder, mason, and leafcutter bees (*Megachilidae*), and two sweat bees that nest in decaying wood, *Lasioglossum cressonii* (Mitchell 1960) and *L. oblongum* (Sakagami & Michener 1962). Above-ground nesting bees included 22 species. The rest of the bees were classified as soil nesting bees, which included 69 species across several groups: (i) long-horned bees (Tribe Eucerini), (ii) mining bees (Andrena spp.), (iii) green bees, (iv) all of the other sweat bees, and (v) any others were classified as soil nesting bees, although natural history observation of many species could not be located.

### Measuring land cover and neighborhood development around study sites

Six-inch resolution digital aerial images were used to classify impervious surfaces such as roads, parking lots, and structures. Unsupervised classification was initiated with 30 classes that were clumped into land cover types. The impervious surface layer from this classification was added to the City of Madison building footprint and road layer to recover impervious surface obscured by tree canopy. Natural vegetation was identified visually within 1000 m of each site and included open canopy, perennial grasses and forbs in greenways, parks, or transportation corridors. Closed canopy forest was also digitized around sites. Each land cover variable was measured as a percent of the landscape, then variables were standardized with a mean of zero and standard deviation of 1 for comparison in analyses. The three land cover types were also consolidated in a distance matrix at each scale. To characterize neighborhood development history, publicly accessible tax assessment data was obtained and property development year was extracted for parcels located within a 200 m radius of each site, from which we extracted an area-weighted average development year for each site. A Bray-Curtis distance matrix was constructed to contrast sites based on the variability of the area-weighted average, median development year, and most recent property development year within the 200m buffer.

### Data analysis

Individual-based rarefaction curves were constructed for each site using the ‘vegan’ R package, and rarefaction-based species richness estimates were compared to observed richness (Oksanen et al. 2018). Rarefied richness did not reach an asymptote, so raw abundance and richness values were used as sampling effort was standardized (Fig. 2). We used linear regression models to test whether land cover and neighborhood development (median property development year) affected bee species richness (α-diversity); separate analyses were conducted for the overall community and three bee guilds. All variables were scaled to a mean of 0 and standard deviation of 1 and top models were selected using stepwise AIC model selection using the ‘MASS’ R package (Ripley et al. 2018). For purposes of comparison we discuss “old” neighborhoods as those with a median development year prior to 1960 and “new” neighborhoods as those with a median development year after 1960. Bee abundance and richness seemed to drop off after this time point, reflecting a qualitative difference rather than a gradual, linear decline. The Moran’s I test was used to check for spatial autocorrelation in model fit for each full and final models, applied using the ‘car’ R package (Fox et al. 2018).

**Figure 2.**
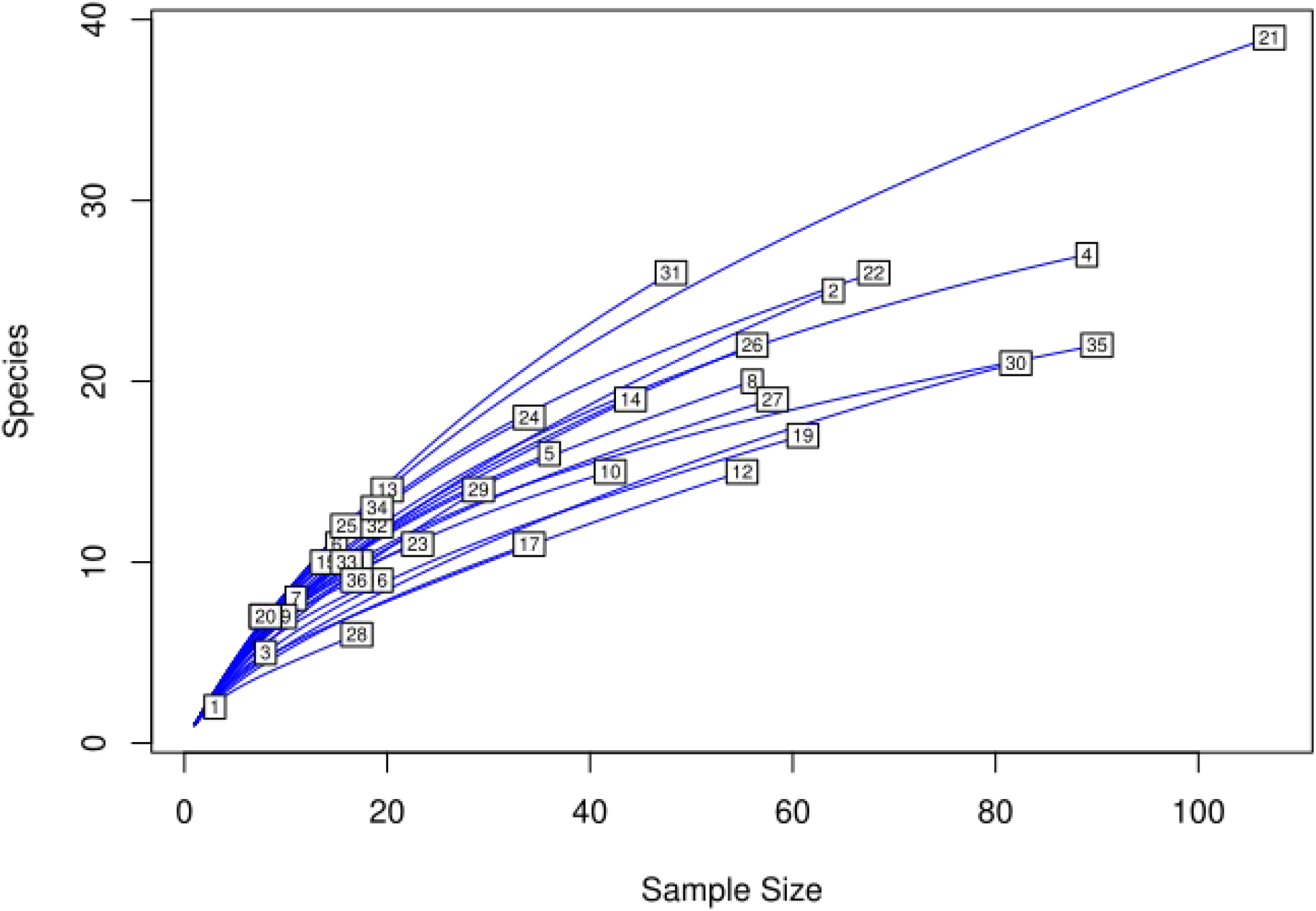
Species accumulation curves for overall site species richness with site numbers.

We used multiple regression on distance matrices (MRM) to assess effects of the various explanatory variables on bee community composition at the landscape scale (β-diversity), which was implemented through the R package ‘ecodist’ (Legendre and Legendre 1998; Goslee and Urban 2017). This allowed us to capture the various multifaceted explanatory variables reflecting heterogeneity of land cover and land use history. MRMs measure the effect magnitude of each explanatory distance matrix using a non-parametric framework and pseudo *t*-tests are used to assess significance of explanatory variables (Goslee and Urban 2017).

## Results

We captured 1331 bees at the 38 sites. Across families, 31% were Apidae, 3% Andrenidae, 55% Halictidae, 8% Megachilidae, and 3% Collitidae. Sites were surrounded by 0 to 43% natural vegetation (Mean = 10.3; SD = 9.1) and 0 to 28% forest (Mean = 5.0; SD = 6.3) with median property development varying between 1920 and 2003 (Mean = 1947; SD = 22). All full bee community and soil nesting bee community analyses were performed across all sites (n = 38). Above ground nesting bee and below ground cavity nesting bee analyses were performed across sites where bees from the nesting guild were present, 32 and 17 sites, respectively.

### Effects of recent property development on bee abundance and diversity

The average bee abundance decreased from 41.7 to 20.8 (*t* = −2.77, df = 25.98, *P* = 0.01), and average species richness from 17.4 to 11.3 (*t* = −2.76, df = 24.16, *P* = 0.01), in pre-1960 compared to post-1960 median development year neighborhoods (Fig. 3). This was driven mainly by decreases in soil-nesting bees, which decreased in abundance from 28.7 to 13.6 bees, and 11.3 to 7.6 species, from old (n = 29) to new (n = 9) neighborhoods. The abundance and richness of above-ground nesting bees, and below-ground cavity nesting bees, were similarly abundant and rich in old and new neighborhoods (*P* > 0.24 for all four metrics). In multiple variate linear regression models including development and land cover variables, a negative influence of recent property development was the strongest predictor of overall bee and soil nesting bee species richness, and the term was included with the negative influence of impervious surface in top models (Tables 1, 2).

**Figure 3.**
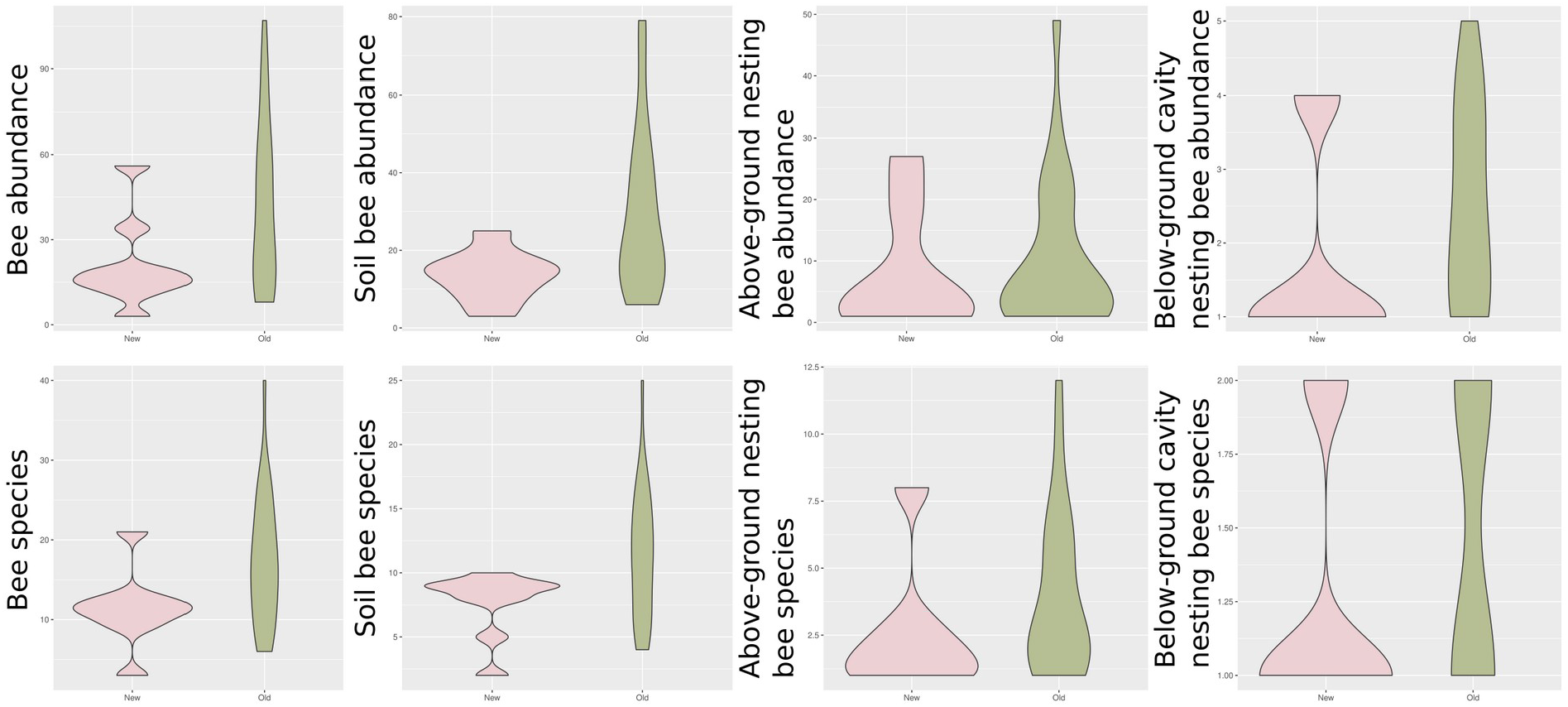
Average bee abundance and species richness for older neighborhoods and recently developed neighborhoods with median property development year in the surrounding 200m landscape sector before or after 1960.

**Table 1.** Results of top AICc-selected multiple linear regression models for species richness of a. the full bee community, b. soil-nesting bees, c. cavity-nesting bees, and d. above-ground bees.

| Full bee community |  |  |  |  |  |
| --- | --- | --- | --- | --- | --- |
| Variable | Estimate | Std Error | *P* | Model *adj R*<sup>2</sup> | *P* |
| Intercept | 16.355 | 1.312 | >0.001 | 0.217 | 0.007 |
| Develop | -5.554 | 2.631 | 0.042 |  |  |
| Imp200 | -2.880 | 1.140 | 0.017 |  |  |
| Soil nesting bee community |  |  |  |  |  |
| Variable | Estimate | Std Error | *P* | Model *adj R*<sup>2</sup> | *P* |
| Intercept | 0.460 | 0.817 | >0.001 | 0.197 | 0.010 |
| Develop | -3.387 | 1.638 | 0.047 |  |  |
| Imp200 | -1.667 | 0.710 | 0.025 |  |  |
| Above ground nesting bee community |  |  |  |  |  |
| Variable | Estimate | Std Error | *P* | Model *adj R*<sup>2</sup> | *P* |
| Intercept | 3.630 | 0.492 | >0.001 | 0.0954 | 0.0475 |
| Imp200 | -1.029 | 0.500 | 0.048 |  |  |
| Below ground cavity nesting bee community |  |  |  |  |  |
| Variable | Estimate | Std Error | *P* | Model *adj R*<sup>2</sup> | *P* |
| Intercept | 0.460 | 0.294 | 0.131 | 0.426 | 0.002 |
| For1000 | -0.357 | 0.494 | 0.477 |  |  |
| For200 | -1.022 | 0.392 | 0.015 |  |  |

**Table 2.** Model average coefficients for the 95% confidence model set of AICc-selected multiple linear regression models for species richness of a. the full bee community, b. soil-nesting bees, c. cavity-nesting bees, and d. above-ground bees.

| Full bee community |  |  |  |  |  |  |  |  |
| --- | --- | --- | --- | --- | --- | --- | --- | --- |
| Variable | Intercept | Forest (1km) | Nat Veg (1km) | Imp (1km) | Forest (200m) | Nat Veg (200m) | Imp (200m) | Development |
| Model Avg Coef | 15.997 | 0.250 | -0.061 | 0.029 | -0.27 | -0.798 | -3.049 | -5.725 |
| Sum of Weights |  | 0.250 | 0.190 | 0.180 | 0.25 | 0.22 | 0.89 | 0.72 |
| Soil nesting bee community |  |  |  |  |  |  |  |  |
| Variable | Intercept | Forest (1km) | Nat Veg (1km) | Imp (1km) | Forest (200m) | Nat Veg (200m) | Imp (200m) | Development |
| Model Avg Coef | 11.134 | 0.807 | 0.329 | -0.131 | -0.327 | -0.457 | -1.724 | -3.608 |
| Sum of Weights |  | 0.290 | 0.200 | 0.180 | 0.21 | 0.22 | 0.82 | 0.73 |
| Above ground bee community |  |  |  |  |  |  |  |  |
| Variable | Intercept | Forest (1km) | Nat Veg (1km) | Imp (1km) | Forest (200m) | Nat Veg (200m) | Imp (200m) | Development |
| Model Avg Coef | 3.695 | -0.169 | -0.641 | -0.054 | -0.009 | -0.337 | -0.679 | -0.316 |
| Sum of Weights |  | 0.180 | 0.360 | 0.180 | 0.18 | 0.22 | 0.67 | 0.27 |
| Below ground cavity bee community |  |  |  |  |  |  |  |  |
| Variable | Intercept | Forest (1km) | Nat Veg (1km) | Imp (1km) | Forest (200m) | Nat Veg (200m) | Imp (200m) | Development |
| Model Avg Coef | 1.434 | -0.129 | 0.092 | -0.027 | 0.368 | 0.081 | -0.092 | -0.017 |
| Sum of Weights |  | 0.520 | 0.170 | 0.200 | 0.51 | 0.15 | 0.14 | 0.12 |

### Effects of surrounding land cover on bee species diversity

In addition to effects of neighborhood development, the proportion of impervious surface also reduced the species richness (α-diversity) of the overall bee community, soil-nesting bees, and above-ground nesting bees (Tables 1, 2). The negative influence of impervious surface on the overall richness of bee species and soil-nesting bees were each about half the magnitude of the property development effect in the scaled regression model. For the overall bee community, the regression model indicates a 2.9 factor decrease in bee species richness per 23% increase in the proportion of impervious surface. The below-ground cavity-nesting bee species richness was negatively associated with surrounding forest cover with a 1.0 factor decrease in bumble bee species with each 12% increase in surrounding forest cover.

### Variation in bee community composition across the study extent

The final multiple regression on distance matrix model (MRM) for the full bee community composition included only the land cover effect (*P* = 0.03) (Table 3). For the soil nesting bee community, there was a clear influence of geographic distance on community dissimilarity (*P* = 0.040) and a land cover effect (*P* = 0.04) (Table 3). The below-ground cavity nesting community composition included a weakly significant influence of geographic distance (P = 0.07) (Table 3). And there were no observed effects of geographic distance or land cover on the above-ground bees (Table 3). None of the bee community final models included significant effects of property development on community composition (Table 3).

**Table 3.** Multiple regression on distance matrices to assess the influence of geographic distance, neighborhood development, and land cover on the full bee community, soil-nesting bees, above-ground nesting bees, and d. below-ground cavity nesting bees.

| Full bee community |  |  |  |  |  |
| --- | --- | --- | --- | --- | --- |
| Variable | Regression Coefficients | P | F-value | Model R<sup>2</sup> | P |
| Intercept | 5.55x10<sup>-1</sup> | 0.81 | 2.78 | 0.02 | 0.007 |
| Geographic distance | 1.03x10<sup>-6</sup> | 0.28 |  |  |  |
| Neighborhood development | 8.61x10<sup>-4</sup> | 0.18 |  |  |  |
| Land cover (200m) | -1.83x10<sup>-4</sup> | 0.04 * |  |  |  |
| Land cover (1000m) | 1.53x10<sup>-4</sup> | 0.43 |  |  |  |
| Full bee community – Final model |  |  |  |  |  |
| Variable | Regression Coefficients | P | F-value | Model R<sup>2</sup> | P |
| Intercept | 0.570 | 0.030 | 9.690 | 0.02 | 0.09 |
| Land cover (200m) | -1.86x10<sup>-4</sup> | 0.03 * |  |  |  |
| Soil nesting bee community |  |  |  |  |  |
| Variable | Regression Coefficients | P | F-value | Model R<sup>2</sup> | P |
| Intercept | 4.92x10<sup>-1</sup> | 1.000 | 4.160 | 0.030 | 0.020 |
| Geographic distance | 2.16x10<sup>-6</sup> | 0.04 * |  |  |  |
| Neighborhood development | 7.51x10<sup>-4</sup> | 0.260 |  |  |  |
| Land cover (200m) | -1.70x10<sup>-4</sup> | 0.04 * |  |  |  |
| Land cover (1000m) | -1.26x10<sup>-4</sup> | 0.500 |  |  |  |
| Soil nesting bee community – Final model |  |  |  |  |  |
| Variable | Regression Coefficients | P | F-value | Model R<sup>2</sup> | P |
| Intercept | 4.91x10<sup>-1</sup> | 0.97 | 7.390 | 0.020 | 0.010 |
| Geographic distance | -1.70x10<sup>-4</sup> | 0.04 * |  |  |  |
| Land cover (200m) | 2.18x10<sup>-6</sup> | 0.04 * |  |  |  |
| Above ground nesting bee community |  |  |  |  |  |
| Variable | Regression Coefficients | P | F-value | Model R<sup>2</sup> | P |
| Intercept | 8.13x10<sup>-1</sup> | 0.110 | 1.590 | 0.01 | 0.36 |
| Geographic distance | -1.84x10<sup>-6</sup> | 0.210 |  |  |  |
| Neighborhood development | 1.65x10<sup>-4</sup> | 0.480 |  |  |  |
| Land cover (200m) | 3.33x10<sup>-4</sup> | 0.680 |  |  |  |
| Land cover (1000m) | -3.71x10<sup>-4</sup> | 0.170 |  |  |  |
| Above ground nesting bee community – Final model |  |  |  |  |  |
| Variable | Regression Coefficients | P | F-value | Model R<sup>2</sup> | P |
| Intercept | 7.69x10<sup>-1</sup> | 0.270 | 2.130 | 0 | 0.14 |
| Land cover (1000m) | -4.00x10<sup>-4</sup> | 0.140 |  |  |  |
| Below ground cavity nesting bee community |  |  |  |  |  |
| Variable | Regression Coefficients | P | F-value | Model R<sup>2</sup> | P |
| Intercept | 3.76x10<sup>-1</sup> | 0.95 | 5.810 | 0.070 | 0.040 |
| Geographic distance | 5.07 | 0.08 . |  |  |  |
| Neighborhood development | -2.55 | 0.63 |  |  |  |
| Land cover (200m) | -4.30x10<sup>-4</sup> | 0.92 |  |  |  |
| Land cover (1000m) | -3.60x10<sup>-3</sup> | 0.6 |  |  |  |
| Below ground cavity nesting bee community – Final model |  |  |  |  |  |
| Variable | Regression Coefficients | P | F-value | Model R<sup>2</sup> | P |
| Intercept | 3.72x10<sup>-1</sup> | 0.96 | 3.810 | 0.030 | 0.070 |
| Land cover (1000m) | 5.23x10<sup>-6</sup> | 0.07 . |  |  |  |

## Discussion

Bees from each nesting guild were observed throughout the City of Madison at both low and high impervious surface sites. This result suggests that in general, bees are able to use small patches of habitat within the most urbanized landscapes of the city (Theodorou 2016; Hall et al. 2016, Daniels et al. 2020). Our observation of an association between impervious surface and reduced bee community richness, especially for soil nesting bees, reflected patterns reminiscent of a 60-year study in Brazil, which documented an increase of impervious surface and decrease in soil bee nests, abundance, and declines of species richness and phylogenetic diversity (Pereira et al. 2020). The negative influence of impervious surface on soil-nesting bees, above-ground cavity-nesting bees, and the entire bee community, may stem from a loss of exposed soil used for nesting habitat, and associated decreases in flowering forbs that bees use as a food resource.

Our finding that more recently developed neighborhoods exhibited lower bee abundance and diversity was not based on our initial expectations of mechanistic associations between land cover transformation and bee habitat provisioning. A negative influence of recent development was observed for the full bee community and soil-nesting bees. While this negative effect may be due to disturbance and soil compaction, we also observed a reduction of structural complexity in recently developed neighborhoods surrounded by more grass lawn and less gardens that may provision diverse types of bee habitat. More established neighborhoods more frequently offered more complex built habitat including rock walls and gardens rather than simple lawn land cover.

While we expected that below-ground cavity nesting bees would be the most impacted by urbanization and impervious surface, we did not observe that result. Bumble bees that comprised this nesting habitat guild can forage long distances, and other studies have observed bumble bee foraging presence to be strongly influenced by floral resources (Turo et al. 2019, Reeher et al. 2020, Cohen et al. 2020) In fact, greater urban cover can sometimes increase the abundance of urban bumble bees in urban gardens, and promote higher in-garden foraging, alongside plant richness as another contributing factor (O’Connell et al. 2020). Another study of urban bumble bees in the American Midwest found that bumble bee abundance and richness were unaffected by the amount of impervious surface across several cities (Reeher et al. 2020).

While geographic distance did not explain the dissimilarity of the full bee community, it contributed to the dissimilarity in soil-nesting and below-ground cavity-nesting bee community composition. This confirmed our hypothesis that generally smaller, soil-nesting bee communities would vary more across the urban study extent. Past studies have confirmed that bee foraging distances are correlated with body size, contributing to patchy distributions of small bee species (Steffan-Dewenter et al. 2002; McKinney 2008). A recent study of pollinators around cotton farms in Texas found no geographic pattern of isolation by distance for bees, but these patterns were observed for beetles and other more movement limited insect taxa (Cusser et al. 2018).

Urbanization can also filter bee community composition, with some evidence that urban bee communities are more homogenous subsets of nearby rural bee communities (Banaszak-Cibicka 2020). In the models for the species composition of the full bee community as well as each nesting habitat guild, property development did not appear to filter the species composition. Surrounding land cover did affect the full bee and soil-nesting bee community dissimilarity. While the influence of land cover significantly influenced the dissimilarity of species assemblages, these factors did not explain much of the variation overall. High species richness of bees was observed across the city, as well as patchy distributions of rare species.

Research documenting responses of bee communities to urbanization is on the rise, but a recent meta-analysis only discovered three published studies assessing the relationships between bee traits and urbanization (Buchholz and Egerer 2020). As urbanization processes continue to transform landscapes around the world, improving our understanding of habitat provisioning and ecosystem services in urban ecosystems is of great importance. Globally, urban bee research is heavily biased towards cities in developed countries with temperate climates (Silva et al. 2021). Improving the targeted nature of urban pollinator research and accomplishing this research in diverse urban landscapes will bolster our capacity to integrate habitat that supplies pollination services and biodiversity to cityscapes around the world.

## Declarations

### Availability of data and material

Bees are submitted to the University of Wisconsin-Madison insect museum.

### Code availability

https://github.com/verawp

### Author contributions

VWP designed study, performed analysis and drafted manuscript, JSB and DWC participated in manuscript writing. All authors gave final approval for publication.

Compliance with ethical standards

---

**Supplemental Table 1.**
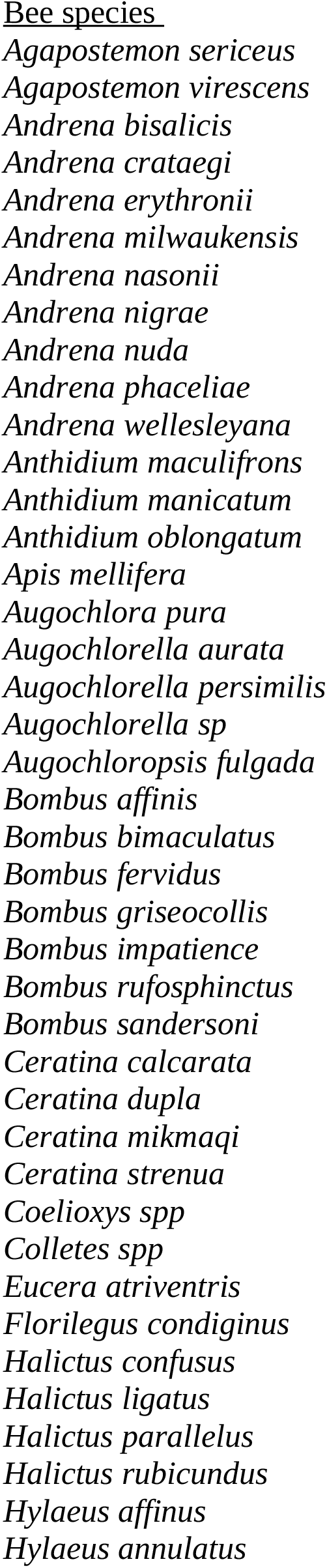
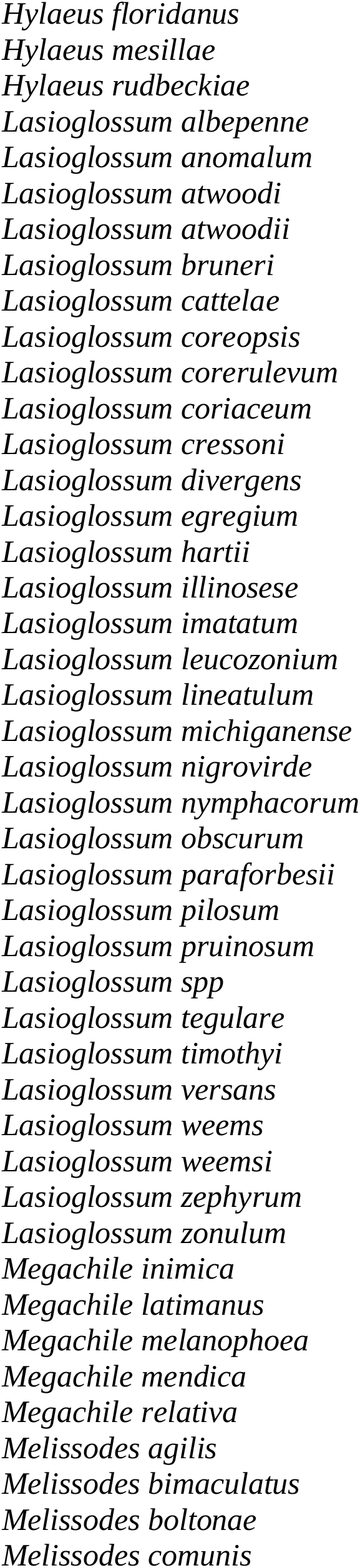
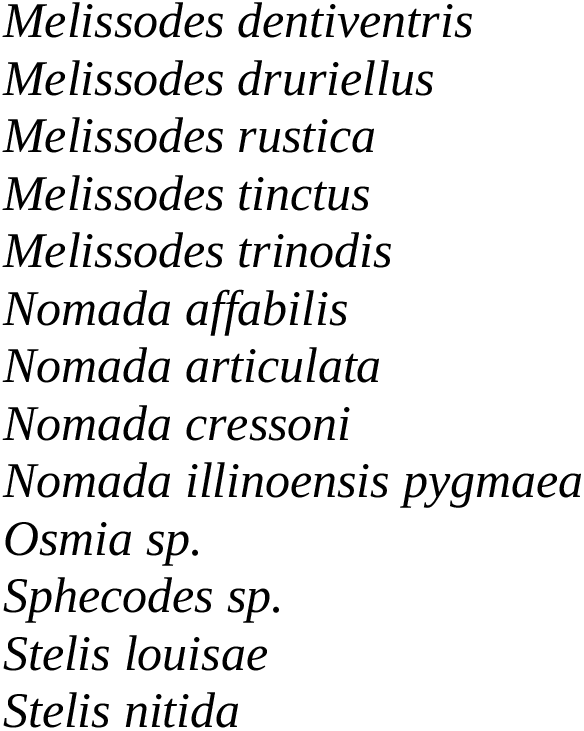

